# Genome-wide mining and comparative analysis of microsatellite markers from *Orientia tsutsugamushi* genomes

**DOI:** 10.1101/2023.02.06.527248

**Authors:** Subhasmita Panda, Subrat Kumar Swain, Basanta Pravas Sahu, Rachita Sarangi

## Abstract

Microsatellite markers, otherwise known as the simple sequence repeats (SSRs), are being used for molecular identification and characterization as well as estimation of evolution pattern of the organism due to their high polymorphic nature. These are tandemly repeated sequences observed almost all organisms and differentially distributed across the genome. Although the primary genome information of *Orientia tsutsugamushi* (OT) suggested the repeats hold the 40% entire of its genome, but lack of characteristic of this repeats increase our interest to study more about it. Thus we investigated a genome-wide presence of microsatellites within nine complete genomes within OT and analyzed their distribution pattern, composition and complexity. The *in-silico* study revealed the genome of OT enrich with microsatellites having a total of 126187 SSR and 10374 cSSR throughout the genome from which 70% and 30% represented within the coding and non coding region respectively. The relative density (RD) and relative abundance (RA) of SSRs were 42-44.43/kb and 6.25-6.59/kb while for cSSRs this value ranged from 7.06-8.1/kb and 0.50-0.55/kb respectively. However, RA and RD were weakly correlate with genome size and incidence microsatellites. The mononucleotide repeats (54.55%) were prevalent over di- (33.22%), tri- (11.88%), tetra- (0.27%), penta- (0.02%), hexanucleotide (0.04%) repeats, with poly (A/T) richness over poly (G/C). Motif composition of cSSRs revealed that maximum cSSRs were made up of two microsatellites having unique duplication pattern such as AT-x-AT, CG-x-CG. More numbers microsatellites represented within the coding region provides an insight into the genome plasticity that may interfere for gene regulation to mitigate with host-pathogen interaction and evolution of the species.

## Introduction

Scrub typhus (ST) is potentially severe but treatable zoonotic bacterial disease caused by a gram negative bacteria, *Orientia tsutsugamushi* (OT) belonging to the genus Orientia of the Rickettsiae family with major genetic changes in peptidoglycan and lipopolysaccharide (LPS) as compared with genus Rickettsia (Xu et al., 2017). OT possesses a 2.1 megabase extremely repetitive single chromosomal genome. Short repetitive sequences, transposable elements (miniature inverted-repeat transposable elements, Group-II intron), and the rickettsial amplified genetic element (RAGE) comprises 42% of the whole genome. The integrase and transposase genes discovered in Integrative and Conjunctive element (ICE) regulate type-IV secretion system as well as potential effector proteins like ankyrin repeat containing protein, histidine kinase, and tetratricopeptide repeats (TPR) domain-containing proteins. The locations of the genes in two complete genome sequences of OT and other rickettsial genomes are scarcely connected due to frequent repeats and mobile DNA elements, and it undergo frequent reshuffling (Salje et al., 2017). OT antigenic heterogeneity hinders the development of broad immunity, allowing for reinfection.

Satellite evolution and biological roles in many species have received a significant interest in research areas (Richard et al., 2008, Garrido-Ramos et al., 2012). Microsatellites are a useful asset for evaluating genetic diversity due to their high mutation rate and ease of experimentation using polymerase chain reaction. These have been widely used for applications such as parentage analysis, population genetics structure, gene mapping and conservation genetics in eukaryotes since 1980s due to their high level of polymorphism (allelic richness), relatively small size, and higher statistical power per locus (rapid analysis protocol) (Wang et al., 2012; Haasl et al., 2010). In recent years the presence of SSRs were well elucidated throughout the genome of viral, bacterial and several other prokaryotic and utilized as a potential tool for strain identification and pathogen evolution (Alam et al., 2013, 2014; Mrázek et al., 2007; Tóth et al., 2000). Within some prokaryotes its polymorphism within coding region regulates the host pathogen interaction as well as recombination that lead to evolution of the species (George et al., 2015). In this study, we extensively examine the presence, size, density, and motif types of several simple and compound microsatellites found in the OT genome. We also demonstrated the correlation between different parameters which influence the distribution of these repeats. The frequency, composition and complexity of various microsatellites in OT genomes may aid in understanding the functional characteristics and host adaptation.

## Materials and method

### Analysis of genomic assembly

At present, around nine OT isolates have been entirely sequenced and their respective size ranged from 1.9Mb-2.4 Mb nucleotides (**Figure-1**). The NCBI database was used to collect all the nine full genome sequences of those isolates for genome-wide *in-silico* microsatellites analysis. Krait v1.0.3 software was used to analyze the distribution of simple and compound microsatellites across the whole genome (Du et al., 2018). We calculated the relative density (RD) and relative abundance (RA) values to compare the genomic sequences of varying lengths. The total length (bp) supplied by each microsatellite per kilobase (kb) of sequence studied denoted by RD, and the number of microsatellite present per kb of the genome denoted by RA. Microsatellites (SSRs, cSSRs) were screened and localized using Krait v1.0.3, as previously described (Qi et al., 2020; Du et al., 2018).

**Figure-1:**
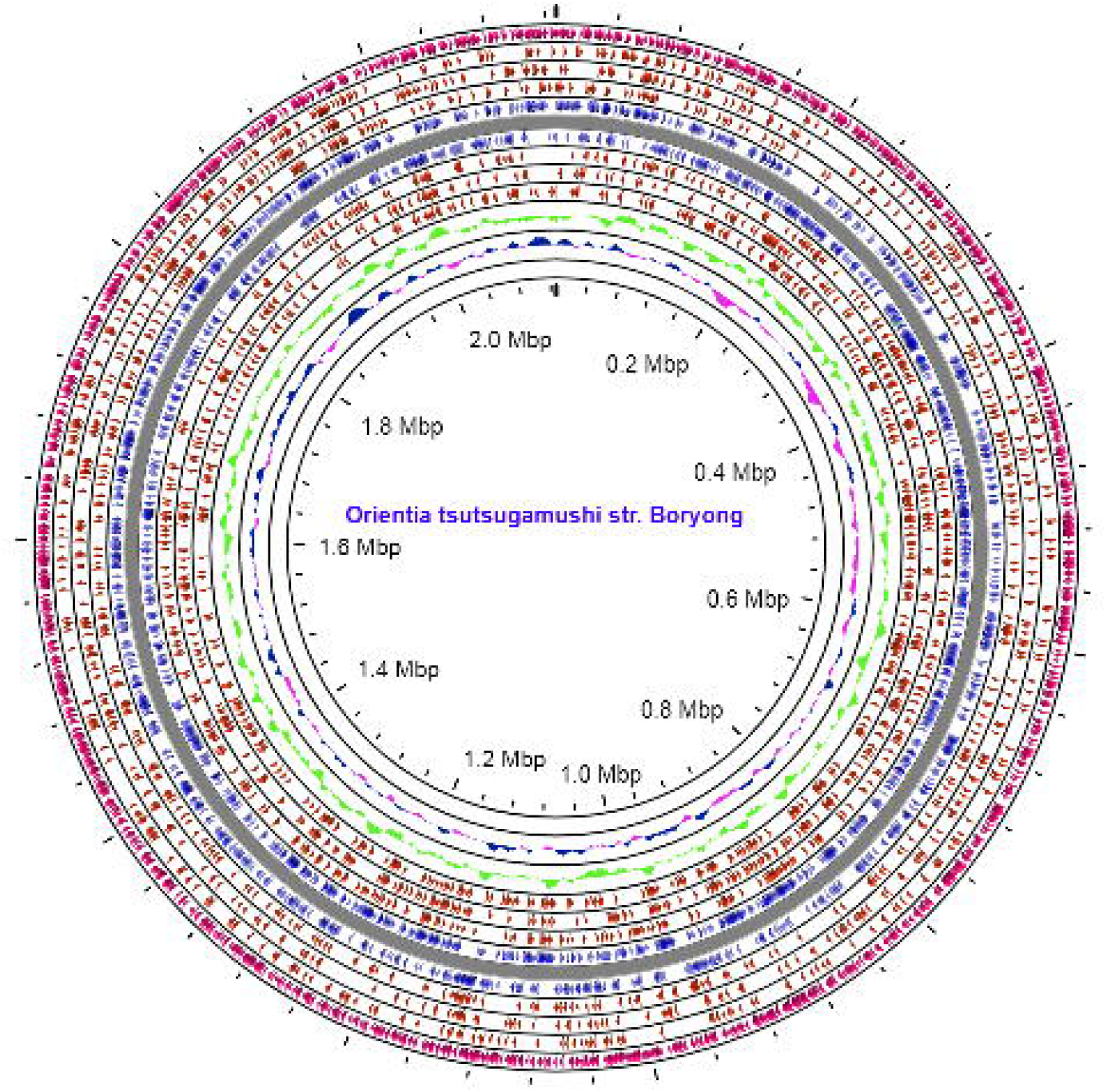
Circos plot representing the genome wide study of SSRs and cSSRs across the nine strains of OT genome. The outer to inner ring demonstrating the SSR, ORF, CDS, repeat region, GC content, GC skew.

### Microsatellite identification

For microsatellite identification the parameters were set as: repetition type: perfect; repeat size: all for mono, di, tri, tetra, penta, and hexa nucleotide, the minimum repeat number is 6, 3, 3, 3, 3, 3. The maximum distance between any two SSRs (dMAX) was 10 bp.

### Statistical analysis

Microsoft Office Excel was used to do the correlation analysis. The Pearson correlation coefficient (R), was determined using GraphPad Prism software v9.4.1 to assess the impact of genome size and GC content on SSRs and cSSRs. A p-value below 0.05 was deemed significant.

## Results

### Occurrence of SSR

Genome wide extraction of microsatellites across nine OT isolates revealed a total of 126187 SSRs and 10374 cSSRs and the incident frequency of SSR per genome ranged from 12335 (str. UT176) to 16150 (str. Karp). The variation in incident frequency may be due to the differential genome size. A highly variable relative abundance (RA) of SSR was observed that ranged from 6.27/kb-6.59/kb, likewise, the relative density (RD) varied from 42 to 44.43/kb (**Figure-2**). Approximately, 70% and 30% of microsatellite motifs were distributed within the coding and non coding region respectively where the functional protein and hypothetical proteins occupied 67% and 3.4% respectively. On examining the SSR units size classes, mono nucleotide repeats were found to be most abundant (54.55%), followed by dinucleotide (33.22%) and trinucleotide (11.88%) in all the genome. The mean of tetra-nucleotide, penta-nucleotide and hexanucleotide repeats were least in number and represented 0.27%, 0.02% and 0.04% within the OT genomes respectively.

**Figure-2:**
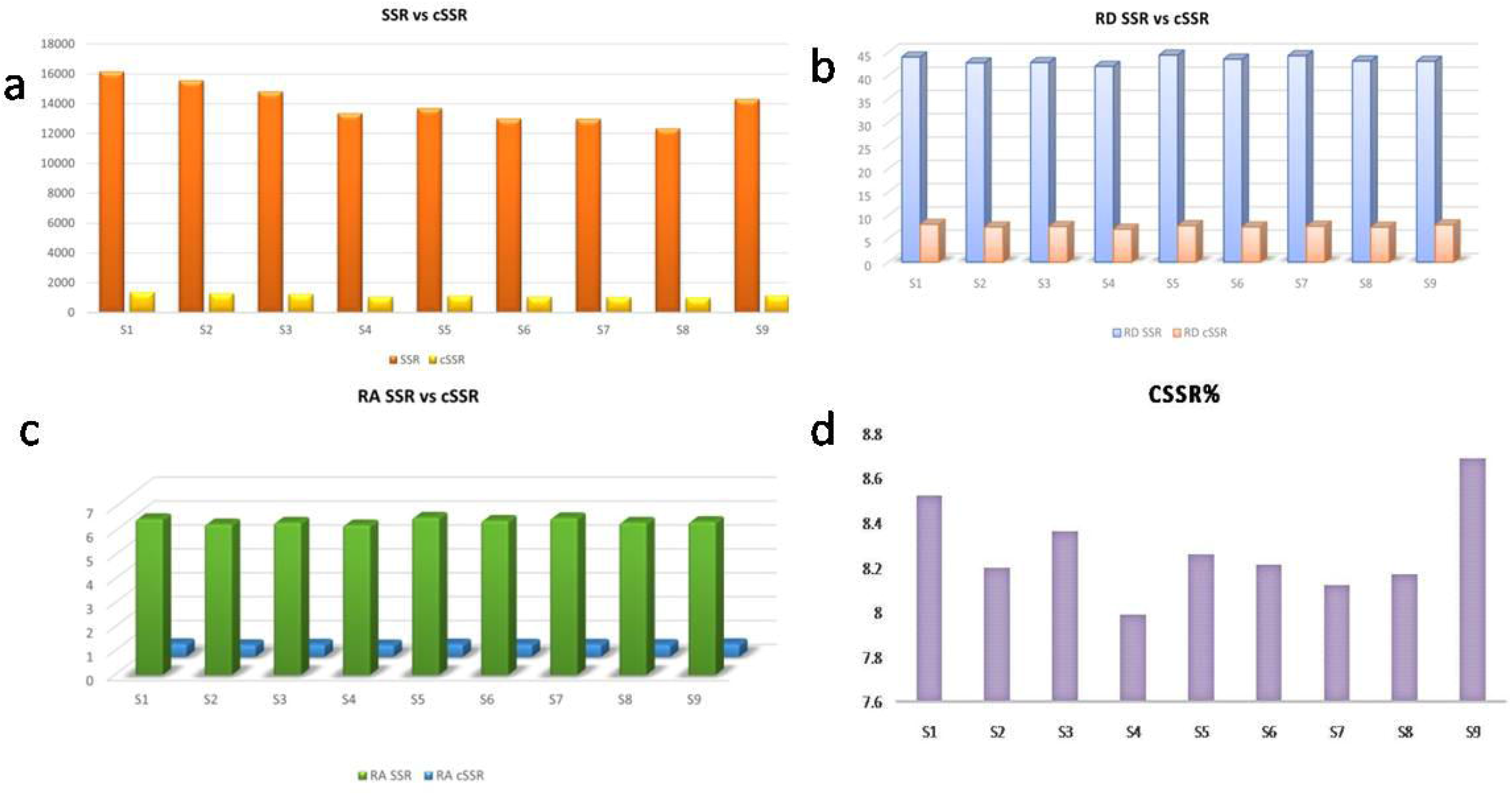
Analysis of SSR and cSSR in OT genome. a) incidence of SSR and cSSR b) relative density (RD) of SSR and cSSR c) Relative abundance (RA) of SSR and cSSR d) % cSSR in OT genome

### cSSR analysis in the OT genome with varying dMAX

The investigation of OT genomes resulted in an observation of a total of 10374 cSSR Karp (OT1) accounted for maximum 1377 cSSR whereas UT176 (OT8) has the minimum 1008 cSSR. The RA and RD ranged from 0.55-0.50 and 7.06-8.1 respectively. The clustering of SSRs could be studied by cSSR incidence and its variability with increasing dMAX to understand if genomes are having SSRs in close vicinity to one another. dMAX is the maximum distance between two SSRs to become a potential cSSR. The value of dMAX can only be set between 10-100 using krait software. To determine the impact of dMAX, the entire nine OT gnomes were analyzed for the number of cSSR with increasing dMAX. We observed an overall increase of cSSR while increased dMAX value for eight genomes, except OT9 which has slightly decrease in dMAX 80 as compared to dMAX70 (**Figure-3**). The percentage of individual microsatellites being part of compound microsatellite (cSSR %) ranged from 8.4-24 (**Table-1**). We observed an overall increase in cSSR% but the same is neither linear nor conforming to any rule. Thus genomes having differential distribution of SSRs might influence the genome diversity and evolution of the OT.

**Table-1:** Overview of microsatellites in *Orientia tsutsugamushi* genome sequence along with other parameters like GC content, incidence of SSR and cSSR with their relative density (RD), relative abundance (RA), %cSSR.

| Sr. no | Acc. No | Names of the strains | Year of strain isolation | Size (bp) | Country | GC content (%) | Total no of SSR | RA | RD | Total no of cSSR | cRA | cRD | % of cSSR |
| --- | --- | --- | --- | --- | --- | --- | --- | --- | --- | --- | --- | --- | --- |
| S1 | NZ_LS398548 | Karp chromosome I | 2018 | 2469803 | United Kingdom | 30.8 | 16150 | 6.53 | 44.01 | 1377 | 0.55 | 8.10 | 8.52 |
| S2 | NZ_LS398551 | Gilliam chromosome I. | 2018 | 2465012 | United Kingdom | 30.5 | 15559 | 6.31 | 42.75 | 1276 | 0.51 | 7.52 | 8.2 |
| S3 | NZ_LS398550 | Kato chromosome I. | 2018 | 2319449 | United Kingdom | 30.8 | 14813 | 6.38 | 42.83 | 1239 | 0.53 | 7.63 | 8.36 |
| S4 | NC_009488 | Boryong | 2007 | 2127051 | South Korea | 30.5 | 13353 | 6.27 | 42.00 | 1067 | 0.50 | 7.06 | 7.99 |
| S5 | NZ_LS398552 | UT76 chromosome I | 2018 | 2078193 | Thailand | 30.5 | 13703 | 6.59 | 44.43 | 1133 | 0.54 | 7.82 | 8.26 |
| S6 | NC_010793 | Ikeda | 2008 | 2008987 | Japan | 30.5 | 12990 | 6.46 | 43.55 | 1067 | 0.53 | 7.55 | 8.21 |
| S7 | NZ_CP044031 | Wuj/2014 chromosome | 2019 | 1972387 | china | 30.5 | 12967 | 6.57 | 44.29 | 1054 | 0.53 | 7.70 | 8.12 |
| S8 | NZ_LS398547 | UT176 chromosome I | 2018 | 1932116 | Thailand | 30.2 | 12335 | 6.38 | 43.16 | 1008 | 0.52 | 7.49 | 8.17 |
| S9 | LS398549.1 | TA686 | 2018 | 2254485 | Thailand | 31 | 14317 | 6.41 | 43.1 | 1153 | 0.55 | 7.99 | 8.69 |

**Figure-3:**
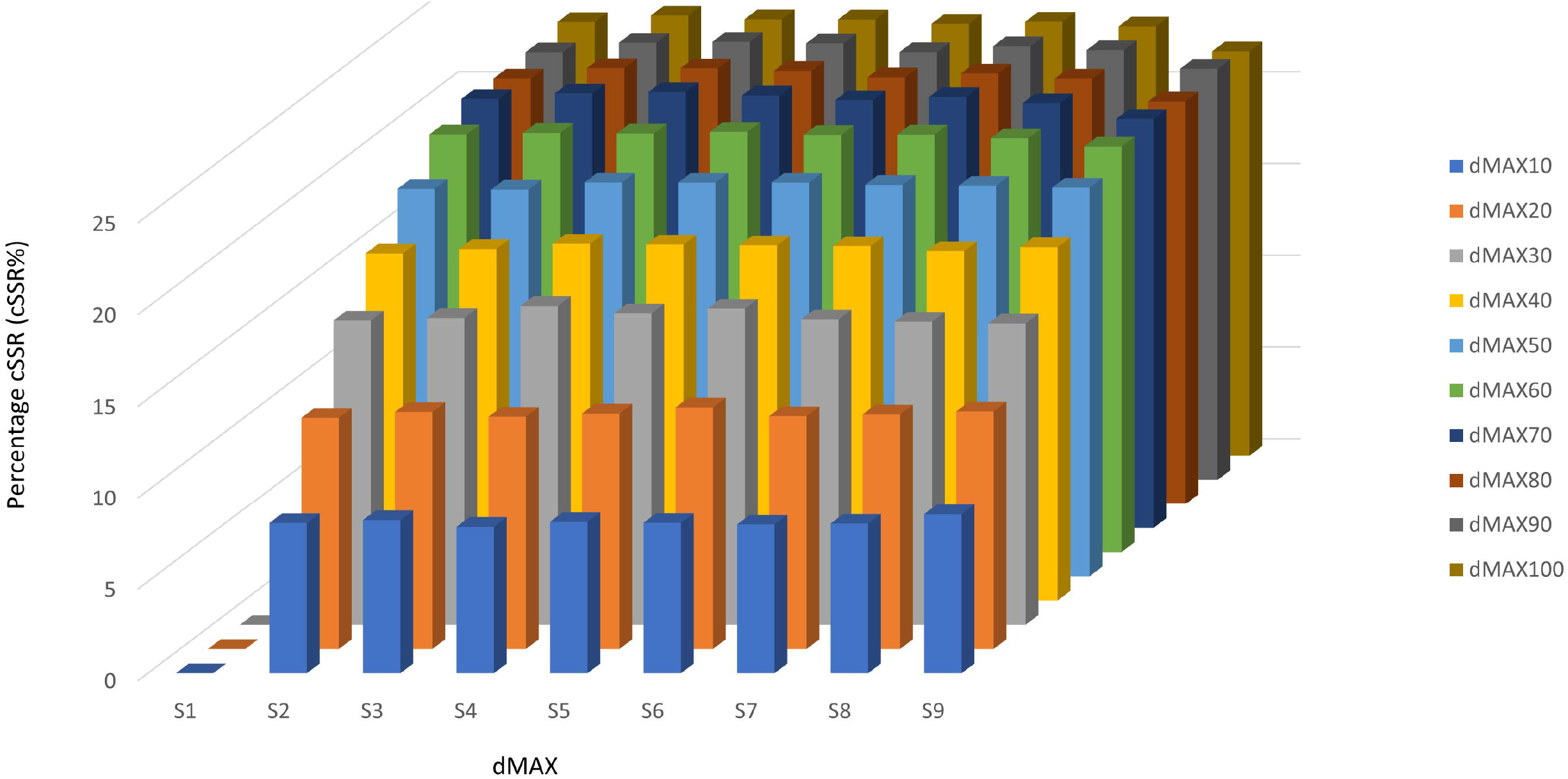
Frequency of cSSR% (percentage of individual microsatellite) by varying the dMAX from 10-100 in nine OT complete genomes. An increased value of cSSR was observed with increase in dMAX.

### Genomic parameters and correlation of SSR and cSSR

We analyzed the possible correlation between the genome size and GC content with the incidence, RA, RD of SSR and cSSR. The genome size of assessed OT has a positive and strong influence on the number of cSSR (r^2^= 0.89; P= 0.001) whereas it has no significant correlation with the GC content (r^2^=0.35; P=0.09). Opposite to that, incidence of SSR has no significant correlation with the genome size (r^2^=0.96; P=2.83) but positively correlated with GC content (r^2^=0.39; P=0.05) of the genome. In contrast, the genome size has no significant correlation with RA (r^2^=0.07; P=0.47) and RD (r^2^=0.04; P=0.56) of SSR and also RA (r^2^=0.02; P=0.66) and RD (r^2^=0.12; P=0.35) cSSR in assessed OT genome. Same as genome size, GC content not correlated with RA (r^2^=3.80; P=0.99) and RD (r^2^=0.002; P=0.96) of SSR and RA (r^2^=0.39; P=0.07) and RD (r^2^=0.35; P=0.09) of cSSRs. The percentage of SSR being the part of cSSR (cSSR %) was positively correlated with the GC content (r^2^=0.67; P=0.006) but not with the genome size (r^2^=0.25; P=0.16) in the OT genome.

### Preferential motif type of SSRs in OT genomes

The divergence of repeat motifs extracted from OT genome ranged from mono to hexa – nucleotide. The prevalent frequency of repeat motifs in each category is a reflection of the GC content of the genome. Most interestingly, poly (A) and poly (T) microsatellite were most prevalent over poly (G) and poly (C) microsatellites as this has been reported as a marker for host determination. This might be attributing to the A/T rich nature of the OT genome. Poly (A) varied from 6 to 18 (OT8), and poly (T) varied from 6 to 22 (OT6). The average frequency of mononucleotide repeats A and T were 3815.77 and 3825.33, the G and C mononucleotide motifs were least represented with the value 8.66 and 10.11 respectively. The dinucleotide repeat motif AT/AT (46.67%) were the most abundant than AT/TA (34.60%), AG/AG (6.36%), AG/TC (6.65%), AG/CT (181.14), AG/GA (4.47%), AC/AC (2.23%), whereas comparative incidence of the least represented CG/CG, CG/GC were 0.23% and 0.81% which was ~58 times less (Figure-4). Tri nucleotide repeats in OT genome reveals ~50 type from which AAT/TAA, AAT/TTA, AAT/ATT, AAT/AAT, AAT/TAT, AAT/ATA, were most abundantly present in OT genome with an average of 10.52%, 10.50%, 9.79%, 8.80%, 8.57%, 8.56% respectively (**Figure-4**). However, TAG, AAG, ATG, GAA, TTG, AGC, ATC and TCA were exhibiting 28.35%, 27.45%, 26.26%, 24.09%, 23.57%, 21.71%, 19.84% and 19.69% respectively. The most common tetra, penta and hexanucleotide repeats are AAAT/AATA, AAGG/GGAA, ATGC/CATG, AATT/TTAA, AAAT/TTAT, AATG/TTAC, AAGGC/CCTTG, AATTG/TAAGT, AAAAT/TAAAA, AGAGC/GAGCA, AAATT/TTAAT, AATGG/TAAGG, AAAGGG/AAGGGA, AAACAG/GTTTCT, ATCATG/ACTAGT, ATATCG/TATAGC, AATTGT/CATTAA, ATCAGT/ACTAGT and AATCGT/GCATTA respectively. It revealed that the frequency of mono, di, and tri repeats varied from each other in different strains of OT.

**Figure-4:**
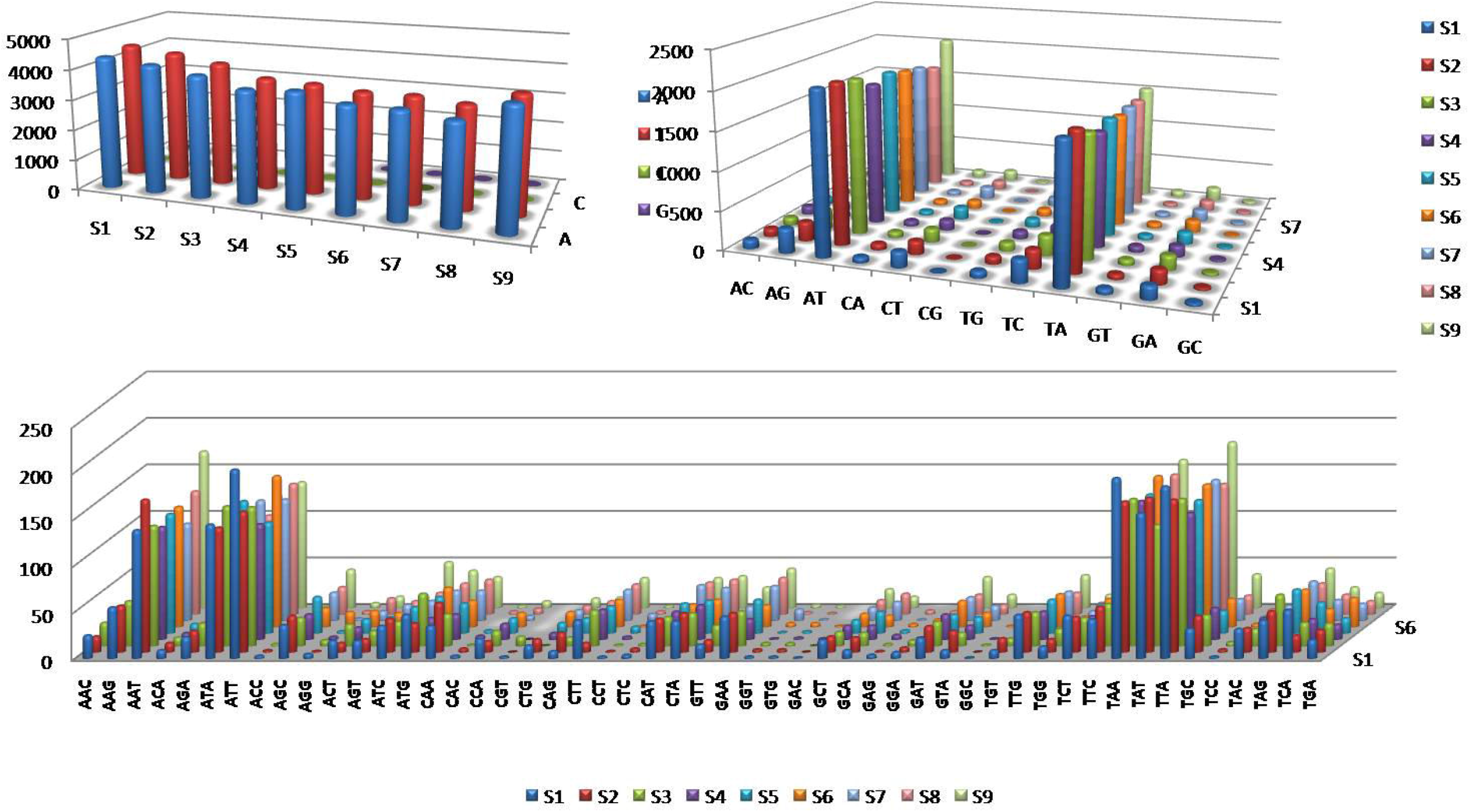
a) Distribution of different motifs of mononucleotide within OT genome b) distribution of different motifs of dinucleotide within OT genome c) distribution of different motifs of trinucleotide within OT genome.

### Motif pattern and complexity of compound microsatellite

Compound microsatellites are composed of two or more adjacent individual microsatellites; therefore, the composition of microsatellites is indeed very complicated due to variable numbers. To counter the effect of differential distribution of compound microsatellite ‘SSR-couples’ term was introduced by Kofler et al., 2008 (Kofler et al., 2008). Motif having the form of [m1]n-xn-[m2]n are known as SSR-couples of motif m1-m2. For example, the compound microsatellite (A)6-x2-(TA)3, (A)6-x4-(TA)3, (A)6-x5-(TA)3 and (A)6-x8-(TA)3 has SSR-couples of motif A-TA. The pattern m1-xn-m2 is considered as ‘2-microsatellite’ and m1-xn-m2-xn-m3 as ‘3-microsatellite’ and so on. In general, the number of compound microsatellite became less with the increase of complexity in the analysis of complete genome of OT. Large cSSRs were observed showing a greater complexity in OT genome than other prokaryotes. Majority of cSSR were composed of two motifs followed by tri, tetra and penta motifs. A number of self-complementary motifs observed in OT genome such as (AT)3-x0-(TA)3, (GA)3-x7-(AG)4, (TA)3-x2-(AT)3, (AT)3-x2-(TA)3, (AT)3-x3-(TA)3, (AT)3-x6-(TA)3, (AT)3-x9-(TA)3, (AT)3-x6-(TA)4, (A)6-x8-(AT)3-x8-(A)6, (A)6-x5-(AT)3-x5-(T)6 which leads to the formation of secondary structure within the genome. The motif composition (CA)n-(X)y-(CA)z forms duplication pattern in which similar motif is located on both ends of the spacer sequence having the motif patter (AT)3-x3-(AT)3, (AT)3-x6-(AT)3, (AT)3-x9-(AT)3, (AT)3-x10-(AT)3, (AT)3-x8-(AT)4. (ATC)3-x3-(ATC)3, (AT)3-x4-(AT)4-x4-(AT)3, (T)10-x6-(T)6, (T)6-x3-(A)6, (TA)3-x4-(TA)3, (TA)3-x5-(TA)3, (TA)3-x8-(TA)3, (A)10-x0-(A)6, (A)6-x1-(A)6, (A)6-x2-(A)7, (A)6-x3-(A)8, (A)6-x4-(A)6, (A)6-x5-(A)9, (A)6-x9-(A)10. We observed the most common microsatellite couples like (TA)-x-(TC), (TG)-x-(TA)-x-(AC), (CA)-x-(GT), (GA)-x-(ATC), (AC)-x-(GT).

## Discussion

Microsatellites are the small motifs of 1-6 base pairs that are tandemly repeated in DNA (Chen et al., 2009; Chen et al., 2010). Although strand-slippage hypothesis are widely employed to explain microsatellite distribution, still they are insufficient to explain the observed divergence of microsatellite dissemination among organism. Microsatellites have been shown to play a key role in transcription, protein function and gene regulation (Kashi, Y. et al., 2007) and utilized as a biomarker for population genetics, linkage association studies (Usdin, K et al., 2008). *Escherichia coli* (E. coli) microsatellites have been extensively studied in prokaryotic genome (Chen et al., 2011), then after the distribution, polymorphism and characterization of microsatellite have been studied in various prokaryotic and eukaryotic genomes, including *Burkholderia pseudomallei* (Ledenyova et al., 2019), *Hemopillus influenza* (Power et al., 2009), *Lactobacillus* (Basharat et al., 2015), *Mycobacterium tuberculosis* (Sreenu et al., 2006), *Mycobacterium bovis* (Sreenu et al., 2006), *Saccharomyces cerevisiae* (Bagshaw A et al., 2008) and DNA viruses such as Human papilloma virus (HPV) (Chen et al., 2012), Adenovirus (Houng et al., 2009), ORF virus (Sahu et al., 2020), Avipoxvirus (Sahu et al., 2022), also in RNA viruses like HIV (Chen et al., 2012), tobamovirus (Alam CM et l., 2013), Carlavirus (Alam CM et al., 2014). Here we investigated the comparative analysis, distribution, and characterization of microsatellites in nine complete OT genomes, which have one of the most complex genomes to date and can serve as an ideal model organism for studying the nature of microsatellites. Several studies have revealed that the highly variable microsatellite loci that exist within genes often encoding for surface antigens in the genomes of pathogenic bacteria are influenced by significant positive selection.

Quantitative comparisons showed that the ranges of SSR and cSSR incidence in the genomes of the OT strains were substantially greater than many bacteria and viruses but smaller than *B. pseudomallei*, which has more SSR and cSSR due to larger genome size. Similar to our study, many studies have been revealed a direct and significant relation between the SSR density and GC content whereas, cSSR incidence was strongly related with the genome size but no impact on RA and RD (Ledenyova et al., 2019). In contrast with that, the number of cSSR weakly correlated with genome size in *E. coli* genome (Chen et al., 2011), this suggests that the association of SSR density with cSSR density may not be dependent on DNA polymerase or replication method suggested by Chen et al., 2011 As compared with many studies the association of densities of cSSR and SSR is dependent on the species and recombination instead of replication. Also no specific distribution pattern of cSSR could be inferred as distribution was not found to be homologous within the genomes. The cSSRs were also not concentrated within specific genes as numerous SSR and cSSRs were present in the hypothetical protein and in the non coding regions so it is difficult to remark on the occurrence of the cSSR with reference to specific proteins or their subunits. Opposite to that Singh et al., 2014 and Sahu et al., 2020 discovered a positive and substantial effect of genome size on RA and RD of both SSR and cSSR in HPV and ORFV genome (Singh AK et al., 2014; Sahu BP et al., 2020). In general the number of compound microsatellites reduces as complexity increases. The variation of cSSR% in different strains of OT was 8-23 and increases with increasing dMAX (10-100) and so as in all studies which suggests that cSSR% is directly related to dMAX (Ledenyova et al., 2019, Chen M et al., 2011, Bagshaw A et al., 2008). In ORFV, 22.1% of cSSR were composed of identical motifs, which were most likely caused by genome duplication. Some research implies that genome duplication may be beneficial to the repeat tendency mechanism (Fadda et al., 2003), which promotes genome size increase in organisms such as yeast (Liti, G. & Louis, E. J 2005; Karaoglu, H et al., 2005).

We accessed the distribution of repeats all over the genome in both the coding and non-coding regions, where the coding sequence correlated with gene arrangement and evolution whereas the non-coding sequence linked with gene regulation and host interaction (Gao et al., 2016). Many researches focused on the influence microsatellites in the coding region and their impact on protein structure and function, as well as codon bias. Furthermore, non-coding sequences have been linked to the host immune response evasion and cellular transformation (Tycowski et al., 2015). In compared to other bacteria and viruses, OT genomes exhibit more SSRs in the coding area than in the non-coding region; this may be owing to the more relaxed selection pressure on the coding region in OT (Sahu et al., 2020). This is the first investigation of microsatellites in the OT genome, and we demonstrated the existence of SSRs across all available whole genomes to date. The increased density of SSR in the coding region emphasizes their importance in gene organization and evolution which needs further in depth evaluation. The fact that cSSRs were conserved across all OT stains highlights their potential use as biomarkers. Kofler et al., 2008 analyzed the distribution of microsatellites across the genome of eight eukaryotic species and hypothesized that nucleotide substitution at the microsatellite tract terminal motifs and the subsequent expansion of the resulting imperfect repeats provides a possible mechanism for compound microsatellite generation. This mechanism for compound microsatellite formation predicts a direct and strong relationship between SSR and cSSR densities (Kofler et al., 2008). To contrary, Chen et al., 2011 investigated the cSSR abundance in the genome of many *E. coli* strains and found only a very weak correlation between SSR and cSSR densities and concluded that microsatellites and cSSR density relationships in eukaryotic and prokaryotic genomes are thought to be influenced by the roles of distinct DNA polymerases and replication mechanism (Chen et al., 2011). cSSRs incidence were strongly correlate with the SSR density which varied dramatically from species to species suggested that the DNA polymerases or replication might not influence the association of SSR and cSSR incidence.

In OT, the analysis of SSRs showed a prevailed of A/T rich di and tri nucleotide repeats rather G/C. Formation of hairpin structure by trinucleotide repeats such as (ATG)n, (ACG)n, (AAG)n, (AGG)n and so on facilitates DNA polymerase slippage during DNA replication which are the main factors of microsatellite instability. Though these are present in OT genome but have low motif copy number suggesting low polymerase slippage which is similar with *B. pseudomallei* (Ledenyova ML et al., 2019). Mononucleotide repeats were preferred over di- and trinucleotide repeats, and their distribution were varied across genomes. Similar to our study, Poly (A/T) repeats were much more abundant than poly (G/C) in HPV genomes whereas in *B. pseudomallei* and *E. coli* G/C were more abundant than A/T. No cSSR contains self complementary motifs observed in OT genome suggested lack of cSSR driven recombination. The prevalence of dinucleotide repeats over trinucleotide repeats maybe linked to the former instability due to a greater slippage rate (Katti et al., 2001), implying that hosts may have a role in the development of dinucleotide repeats within OT genome.

## Conclusion

In conclusion, the study observed the imperfect character and low copy number of motifs that is driven from the investigation of incidence, composition, and complexity of different microsatellites in OT genome. Despite the belief that host adaptation is the primary factor in microsatellite variation, evidence suggests that microsatellite composition is species-specific within the genome rather than host-specific. The study of microsatellites in OT genome is a key step towards better understanding the nature, function and evolutionary analysis of the species. This result may be consider as useful tool for strain identification, diversity estimation and multiple genome analysis have revealed non-intuitive subtleties in the population structure concerning the distribution of SSRs across the OT pan-genome. Because of the genome diversity, OT interacts differently with host-cellular mechanism and the interactions may have significant effect on clinical severity of the diseases.

## Declaration

## Acknowledgement

The authors acknowledge the Medical Research Laboratory of Institute of Medical Sciences and SUM hospital for providing the lab facility, Siksha ‘O’ Anusandhana (deemed to be) University and to SOA University for providing the Ph.D. fellowship as financial support.

## Author Contribution statement

SP and SKS did the bioinformatics analysis, and written the manuscript. SP and SKS constructed the figures. BPS has generated the circos plot. RS has designed the study; BPS executed the study and reviewed the manuscript.

